# A pleiotropic role for FGF signaling in mammary gland stromal fibroblasts

**DOI:** 10.1101/565267

**Authors:** Zuzana Koledova, Jakub Sumbal

## Abstract

Fibroblast growth factor (FGF) signaling is crucial for mammary gland development. While multiple roles for FGF signaling in the epithelium were described, the function of FGF signaling in mammary stroma has not been elucidated. In this study, we investigated FGF signaling in mammary fibroblasts. We found that mammary fibroblasts express FGF receptors 1 and 2 and respond to FGF ligands. In particular, FGF2 and FGF9 induce sustained ERK1/2 signaling and promote fibroblast proliferation and migration in 2D. Intriguingly, only FGF2 induces fibroblast migration in 3D extracellular matrix (ECM) through regulation of actomyosin cytoskeleton and promotes force-mediated collagen remodeling by mammary fibroblasts. Moreover, FGF2 regulates production of ECM proteins by mammary fibroblasts, including collagens, fibronectin, osteopontin, and matrix metalloproteinases. Finally, we show that FGF2 signaling in mammary fibroblasts enhances fibroblast-induced branching of mammary epithelium. Our results demonstrate a pleiotropic role for FGF signaling in mammary fibroblasts with implications for regulation of mammary stromal functions and epithelial branching morphogenesis.

**Summary statement:** FGF signaling in mammary fibroblasts regulates fibroblast proliferation, migration, extracellular matrix production and remodeling, and fibroblast-mediated mammary epithelial branching morphogenesis.

## Introduction

FGF signaling is a crucial pathway that regulates vertebrate development from the earliest embryonic stages throughout lifetime (Turner and Grose, 2010). Importantly, FGF signaling has a conserved role in regulation of branching morphogenesis, governing development of branched organs such as fly trachea and mammalian lung, salivary gland, kidney and mammary gland (Affolter et al., 2009; Lu and Werb, 2008). In mammals, FGF signaling comprises 22 FGF ligands, 18 of which act through 4 transmembrane tyrosine kinase receptors (FGFR1-4). Ligand-binding specificity of FGFR1-3 is generated by alternative splicing of the extracellular immunoglobulin domain III, creating IIIb and IIIc variants of FGFR1-3. Binding of FGF ligand to FGFR requires co-factor (heparan sulfate) and results in receptor dimerization, phosphorylation and activation of downstream signaling pathways, including Ras-MEK-ERK, PI3K-AKT, PLCγ and STAT3 signaling pathways (Turner and Grose, 2010).

FGF signaling is essential for normal mammary gland development. Loss of *Fgf10* or its receptor *Fgfr2* results in a failure to form mammary placodes during embryogenesis (Kim et al., 2013; Mailleux et al., 2002). Conditional deletion of epithelial *Fgfr1* or *Fgfr2* results in transient developmental defects in branching morphogenesis (Lu et al., 2008; Parsa et al., 2008; Pond et al., 2013), while simultaneous deletion of both *Fgfr1* and *Fgfr2* in the mammary epithelium compromises mammary stem cell activities (Pond et al., 2013). Several FGF ligands are produced by the pubertal mammary gland stroma, including FGF2, FGF7, FGF9, and FGF10, and regulate distinct aspects of epithelial morphogenesis (Zhang et al., 2014b). However, the role of FGF signaling in mammary stroma has not been elucidated.

Mammary stroma consists of extracellular matrix (ECM) and several cell types, including fibroblasts, adipocytes, immune and endothelial cells, and has an instructive role in regulating mammary gland development (Nelson and Bissell, 2006; Wiseman and Werb, 2002). Through providing paracrine signals, mechanical cues and organization of 3D ECM scaffold, mammary stroma guides mammary epithelial branching morphogenesis from embryonic stage throughout the puberty, epithelial differentiation to milk-producing alveoli during lactation, and epithelial remodeling during involution (Polyak and Kalluri, 2010; Schedin and Hovey, 2010).

The important role of fibroblasts in modulating mammary epithelial response was described more than 30 years ago (Haslam, 1986), yet the mechanisms of action of fibroblasts in mammary gland development have only recently started to be revealed. In three-dimensional (3D) co-cultures in collagen, fibroblasts induced formation of invasive branched mammary epithelial structures through paracrine hepatocyte growth factor signaling (Zhang et al., 2002). Fibroblasts were found essential for formation of mammary epithelial ductal structures upon transplantation of human mammary organoids into humanized mouse fat pads (Kuperwasser et al., 2004) and in 3D co-culture of MCF10A cells (Krause et al., 2008). Genetic mouse models demonstrated critical roles of fibroblast-mediated paracrine signaling and ECM remodeling in mammary gland development (Hammer et al., 2017; Jones et al., 2019; Koledova et al., 2016; Peuhu et al., 2017) and revealed regulation of these functions by receptor tyrosine kinase signaling, including PDGFR and EGFR-ERK1/2 pathways (Hammer et al., 2017; Koledova et al., 2016).

In this study, we investigated FGF signaling in mammary fibroblasts and its implications for mammary gland development using 2D and 3D cultures of primary mammary fibroblasts. We present our findings on the components and activity of FGF signaling in mammary fibroblasts, and the roles of FGF signaling in regulation of fibroblast proliferation, migration, ECM production and remodeling, and fibroblast-mediated mammary epithelial branching morphogenesis.

## Results

### FGFR1 and FGFR2 are expressed in mammary fibroblasts

To investigate the FGF signaling machinery in mammary fibroblasts, we isolated fibroblasts as well as epithelial organoids from mammary glands of pubertal mice and we analyzed expression of FGFR variants (*Fgfr1-4* genes and their variants IIIb and IIIc, further referred to as “b” or “c”) in these cell populations by qPCR. The purity of cell populations was checked by expression of epithelial marker *Cdh1* and mesenchymal marker *Vim* (Fig. 1A). We detected expression of a wide range of *Fgfr* genes in mammary epithelium, including *Fgfr1b, Fgfr1c, Fgfr2b, Fgfr3b, Fgfr3c, Fgfr4*. However, in mammary fibroblasts, only *Fgfr1c* and *Fgfr2c* were expressed (Fig. 1A).

**Figure 1.**
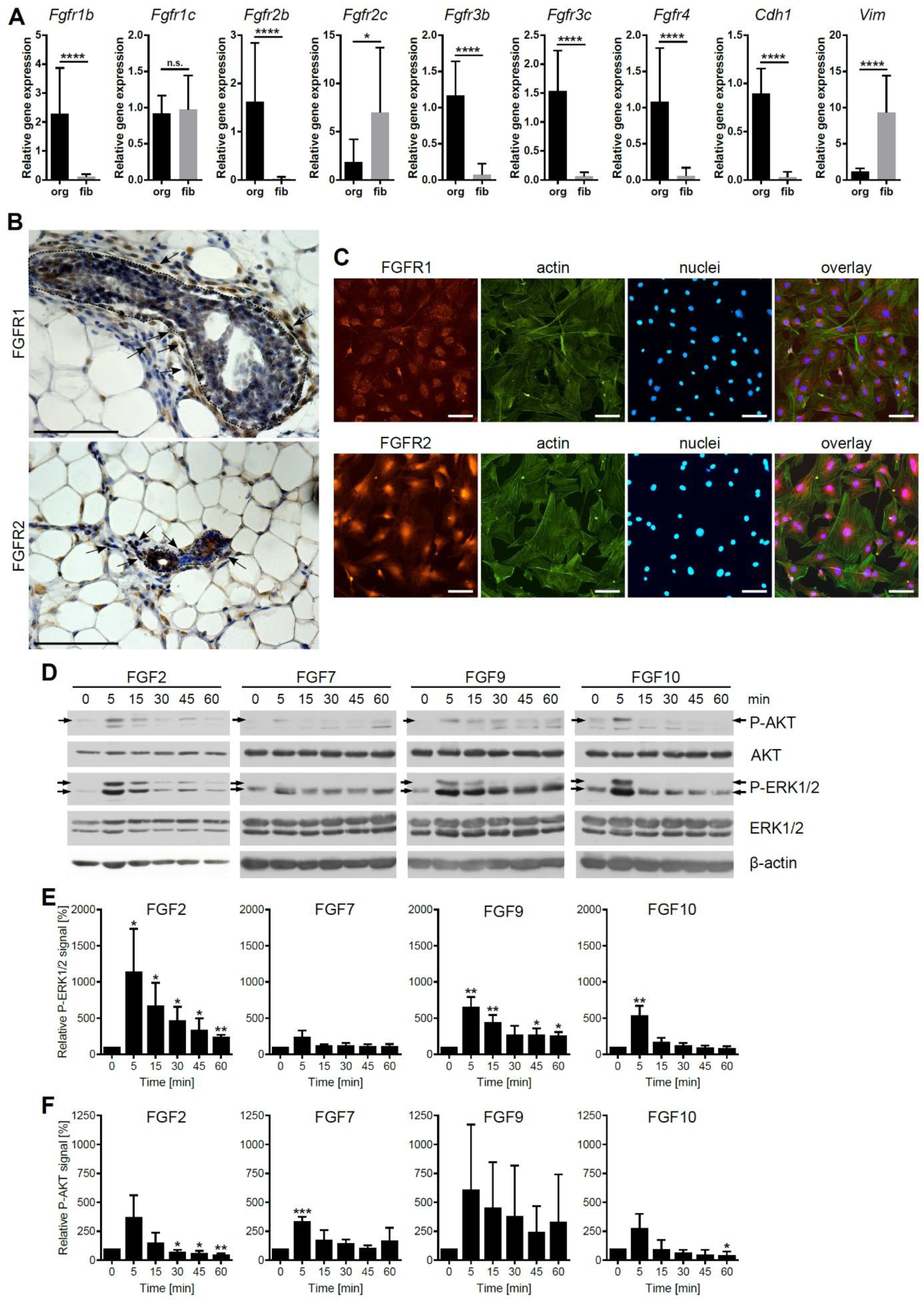
Mammary fibroblasts express FGFR1 and FGFR2 and respond to FGF ligands. (**A-C**) Analysis of FGFR expression in mammary fibroblasts. (**A**) Relative expression of *Fgfr1-4* gene variants in mammary fibroblasts, compared to mammary epithelial organoids. Data are presented as mean + s.d., n = 9 organoid and 22 fibroblast samples. *P < 0.05; ****P < 0.0001 (unpaired Student’s t-test). (**B**) Detection of FGFR1 and FGFR2 in whole mount mammary gland by immunohistochemistry. Dotted lines encircle mammary epithelium. The arrows indicate some of the fibroblasts. Notice expression of both FGFR1 and FGFR2 in mammary fibroblasts as well as epithelium. Blue indicates nuclei stained by hematoxylin. Scale bar, 100 μm. (**C**) Detection of FGFR1 and FGFR2 in primary mammary fibroblasts by immunofluorescence. Actin was stained with phalloidin, nuclei were stained with DAPI. Scale bar, 100 μm. (**D-F**) Western blot investigation of activation of ERK1/2 and AKT signaling pathways in response to FGF ligands. (**E, F**) Quantification of (E) P-ERK1/2 signal, normalized to ERK1/2, or (F) P-AKT signal, normalized to AKT. The bars represent mean + s.d., n = 3. The stars indicate significant change in comparison to cells not treated with FGF (time point 0 min), *P < 0.05; **P < 0.01; ***P < 0.001 (Student’s t-test).

To test expression of FGFR1 and FGFR2 in fibroblasts on protein level, we performed immunohistochemical staining of mammary gland tissue (Fig. 1B) and immunofluorescent staining of mammary fibroblasts in culture (Fig. 1C). Fibroblasts in mammary gland tissue sections, identified as spindle shaped-cells surrounding the epithelium, expressed both FGFR1 and FGFR2 (Fig. 1B). Similarly, primary fibroblasts cultured in vitro, positive for fibroblast markers platelet-derived growth factor receptor alpha (PDGFRα) and vimentin (Fig. S1), expressed both FGFR1 and FGFR2. FGFR1 was localized mostly to the membrane and cytoplasm, while FGFR2 appeared predominantly localized to the nucleus or perinuclear region (Fig. 1B, C).

### FGF ligands induce activation of intracellular signaling pathways downstream of FGFR in mammary fibroblasts

Next we tested four FGF ligands – FGF2, FGF7, FGF9, and FGF10 – which were reported to be expressed in pubertal mammary gland (Zhang et al., 2014b), for their ability to induce FGFR signaling in mammary fibroblasts. To this end, we investigated activation of main signaling pathways downstream of FGFR, ERK1/2 and AKT, by Western blot detection of activated (phosphorylated) forms of ERK1/2 and AKT after addition of FGF ligands to serum-starved fibroblasts. We found that all four FGF ligands induced phosphorylation of ERK1/2 and AKT 5 min after FGF treatment (Fig. 1D-F). However, only in response to FGF2 or FGF9, ERK1/2 phosphorylation (and AKT phosphorylation, too, in the case of FGF2) was sustained beyond 5 min and displayed typical dynamics with signaling maximum at 5 min, followed by gradual decrease of phosphorylation during 60 min after FGF treatment. Activation of ERK1/2 and AKT after FGF7 or FGF10 treatment was only transient and diminished within 15 min after FGF treatment. Therefore, FGF7 and FGF10 were unlikely to induce cellular response.

### FGF2 and FGF9 regulate mammary fibroblast proliferation and migration

In many cell types, FGF signaling is an important regulator of cell proliferation and survival. Therefore, we tested the importance of FGF signaling in mammary fibroblast proliferation and survival using FGFR inhibitor BGJ398 (Guagnano et al., 2011) and SU5402 (Sun et al., 1999). We found that both FGFR inhibitors efficiently inhibit fibroblast proliferation/survival in a dose-as well as cell density-dependent manner (Fig. S2A, B). The half maximal inhibitory concentration (IC_50_) of BGJ398 was 6.98 nM for less confluent cells, and 11.41 nM for more confluent cells. The IC_50_ of SU5402 was 284 nM and 1110 nM for less and more confluent cells, respectively.

Encouraged by the results from FGFR inhibitor assays that suggested that FGF signaling plays a role in fibroblast proliferation and survival, we tested FGF ligands for their effects on fibroblasts. Using MTT proliferation assay, we found that FGF2 and FGF9 enhanced fibroblast proliferation, while FGF7 and FGF10 did not (Fig. 2A, B).

**Figure 2.**
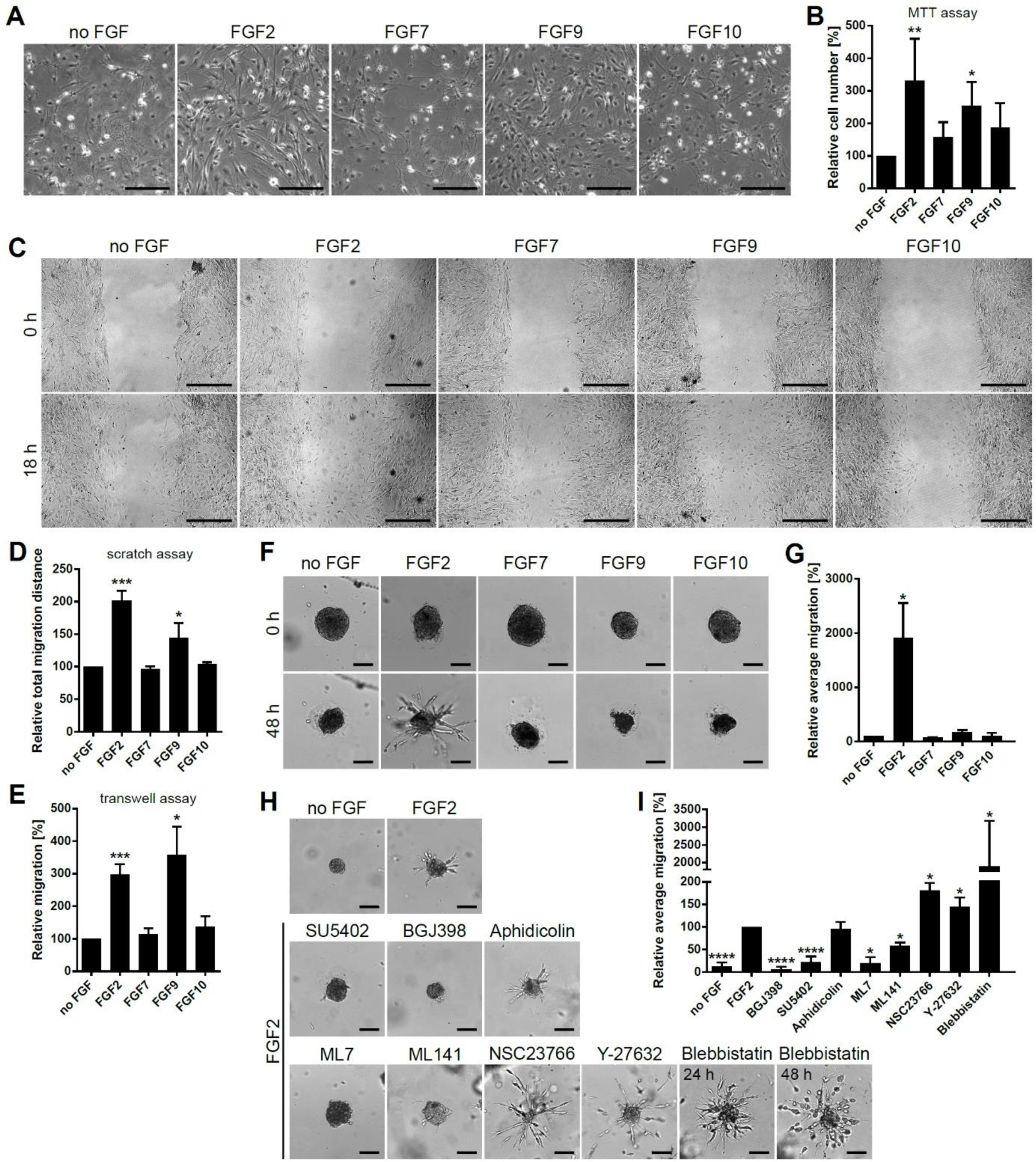
FGF signaling regulates fibroblast proliferation and migration. (**A, B**) FGF proliferation assay. (**A**) Representative photographs of fibroblasts treated with FGF ligands for 24 h, as indicated. (**B**) MTT proliferation assay. The bars represent mean + s.d., n = 5-9. The stars indicate significant change in comparison to cell with no FGF. *P < 0.05; **P < 0.01 (ratio paired Student’s t-test). (**C, D**) Scratch assay. (**C**) Representative photographs of scratch areas at the beginning (0 h) and end of the assay (18 h after scratch). Scale bar, 100 μm. (**D**) Quantification of scratch assay. The bars represent mean + s.d., n = 2-4. The stars indicate significant change in comparison to cell with no FGF. *P<0.05; ***P<0.001 (unpaired Student’s t-test). (**E**) Transwell migration assay. The plot shows mean + s.d., n = 3-7. *P<0.05; ***P<0.001 (ratio paired Student’s t-test). (**F-I**) Fibrosphere 3D migration assay. The fibrospheres were embedded in 3D Matrigel, treated with FGF ligands and/or inhibitors as indicated and imaged by time-lapse microscopy. (**F, G**) FGF2 induces fibroblast migration in Matrigel. (**H, I**) FGF2-induced migration is susceptible to inhibitors of FGFR (0.2 μM BGJ398, 10 μM SU5402), MLCK (5 μM ML7) or Cdc42 inhibitor (5 μM ML141), and potentiated by inhibitors of ROCK (5 μM Y-27632), Rac1 (50 μM NSC23766) or myosin II (25 μM Blebbistatin). (**F, H**) Representative photographs of fibrospheres at 0 h and 48 h (**F**), or at 48 h (**H**). Scale bar, 50 μm. (**G, I**) Quantification of fibroblast migration in 3D Matrigel. The plots show average migration as mean + s.d., n = 2-4. *P < 0.05; ****P < 0.0001 (unpaired Student’s t-test).

Next we tested the effect of FGF ligands on fibroblast migration in 2D using two different assays: scratch assay (or wound healing assay) and transwell migration assay. In the scratch assay, fibroblasts were cultured to confluence, then the cell covered area was scratched to remove cells and to create a cell-free zone, into which the cells can migrate by non-directional migration. We found that fibroblasts migrated into the scratch area even without any FGF in the medium. FGF2 and FGF9 significantly enhanced fibroblasts migration, while FGF7 or FGF10 did not increase fibroblast migration over the baseline (Fig. 2C, D). In transwell migration assay, we tested the capacity of different FGF ligands to induce directed fibroblast migration: from one side of porous membrane in medium with no FGF (upper well), to the other side of the membrane with medium supplemented with different FGF ligands (bottom well). We observed increased migration of fibroblasts towards the medium with FGF2 and FGF9 (Fig. 2E). FGF7 or FGF10 did not act as chemoattractants for the fibroblasts (Fig. 2E).

### FGF2 regulates mammary fibroblast migration in 3D ECM

Subsequently we tested the ability of FGF ligands to induce fibroblast migration in 3D ECM. We employed a fibrosphere 3D migration assay: Fibroblasts were aggregated into a spheroid (fibrosphere), embedded in 3D Matrigel and treated with different FGF ligands. We found that FGF2 was the only of the FGF ligands tested that effectively induced fibroblast migration in 3D Matrigel, visible as radial protrusions of spindle-shaped cells coming out from the fibrosphere in streaks (Fig. 2F, G).

We analyzed the mechanism of FGF2-induced 3D migration more deeply using specific inhibitors. We confirmed that it was FGFR dependent because FGFR inhibitors BGJ398 and SU5402 efficiently abrogated it (Fig. 2H, I). We also found out that the migration in 3D did not require fibroblast proliferation because DNA polymerase inhibitor aphidicolin did not affect it (Fig. 2H, I). Because contraction of actomyosin cytoskeleton is crucial for cell migration, we tested the requirement for several proteins involved in regulation of cytoskeleton contractility. We observed that FGF2-induced 3D migration was efficiently inhibited by Rac1 inhibitor (NSC-23766), Cdc42 inhibitor (ML141) or myosin light chain kinase (MLCK) inhibitor (ML7) but enhanced by Rho-associated protein kinase (ROCK) inhibitor (Y-27632) (Fig. 2J, K). Interestingly, blebbistatin, an inhibitor of myosin II, increased cell migration in 3D ECM and it also changed the mode of cell migration to a more amoeboid-like migration, characterized by a rounded cell body with thin protrusions and decreased migration in streaks (Fig. 2J, K).

### FGF2 promotes collagen remodeling by mammary fibroblasts

During mammary branching morphogenesis, the pattern of collagen fibers in ECM helps to guide mammary ductal outgrowth (Brownfield et al., 2013; Hovey et al., 2002; Ingman et al., 2006; Peuhu et al., 2017). Fibroblasts generate traction forces and reorganize collagen fibers through interaction with ECM and cell migration in response to mechanical and chemical cues (Grinnell and Petroll, 2010; Tschumperlin, 2013). Because our results demonstrated the role for FGF2 in fibroblast migration in 3D ECM (Fig. 2F-K), we investigated the role of FGF signaling in force-mediated collagen remodeling by mammary fibroblasts using collagen contraction assay. Mammary fibroblasts were embedded in floating collagen gels and cultured with FGF ligands or serum and the extent of collagen remodeling was assessed by the decrease of collagen gel size. The gels with fibroblasts that were not exposed to any FGF showed small contraction, while the gels treated with serum contracted to the highest extent (Fig. 3 A, B). FGF2 induced significant collagen gel contraction, while FGF7, FGF9, or FGF10 did not (Fig. 3 A, B). FGF2-induced gel contraction was abrogated by FGFR inhibitors BGJ398 or SU5402 but it was refractory to aphidicolin, an inhibitor of cell proliferation (Fig. 3 A, C). Inhibitors PP2, NSC23766, Y-27632, blebbistatin, or a combination of Rhosin and Y16 inhibited FGF2-induced gel contraction, demonstrating that generation of mechanical force efficient to reorganize collagen required Src, Rac1, ROCK, myosin II, and RhoA, respectively. Inhibitors of matrix metalloproteinases (MMPs) or lysyl oxidase (LOX) (GM6001 and BAPN, respectively) did not abrogate FGF2-induced collagen gel contraction (Fig. 3 A, C).

**Figure 3.**
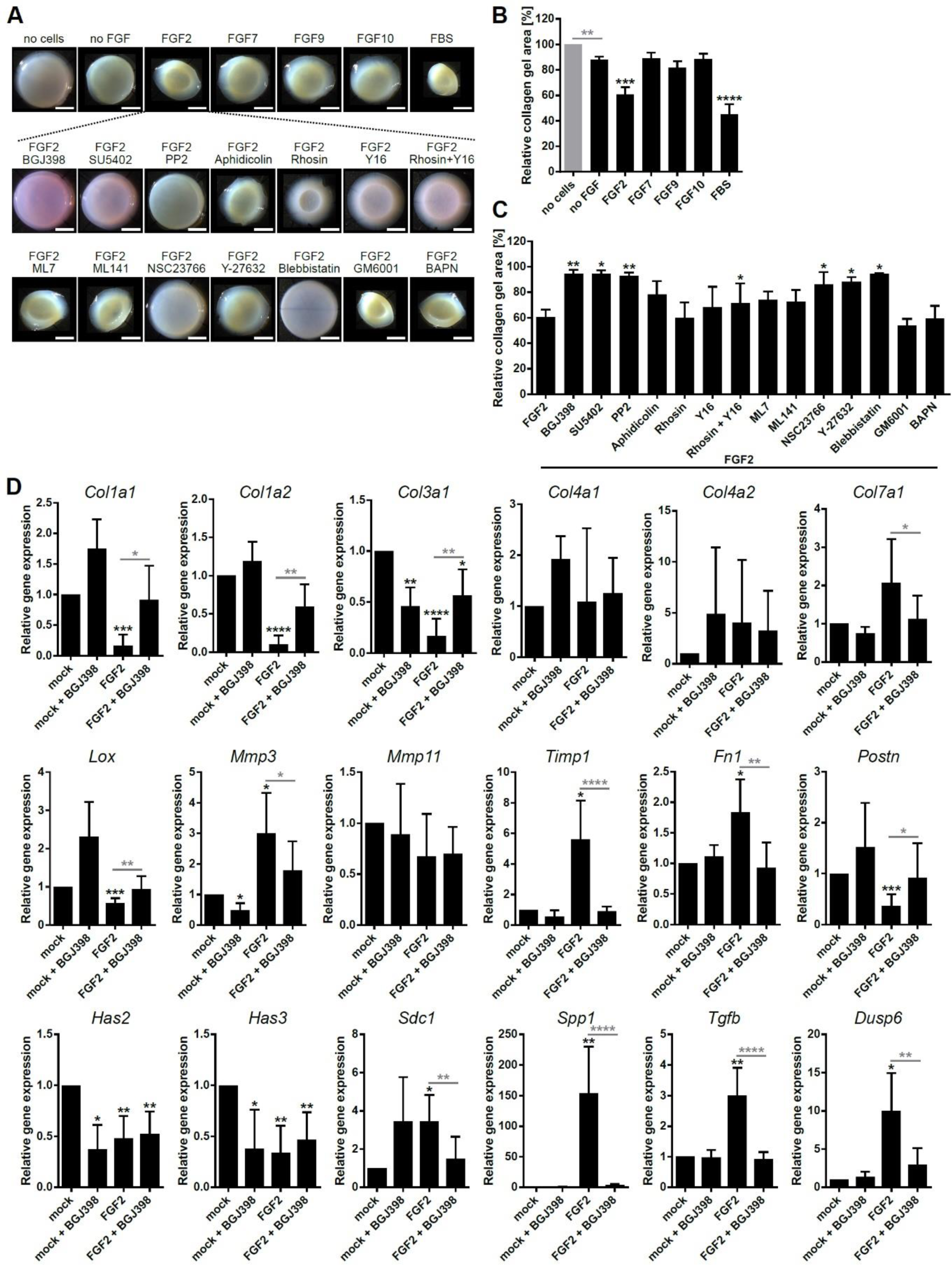
FGF2 regulates ECM organization and synthesis. (**A-C**) Collagen contraction assay. (**A**) Representative photographs of collagen gels. Scale bar, 5 mm. (**B, C**) Quantification of collagen gel contraction in response to (**B**) FGF ligands or serum, or (**C**) FGF2 and inhibitors as indicated. The plots show mean + s.d., n = 5-11. **P < 0.01; ****P < 0.0001 (Student’s t-test). (**D**) FGF2 regulates expression of ECM genes. The plots show relative expression of candidate genes on day 3 of FGF2 treatment as mean + s.d., n = 3. Black stars indicate significant change compared to mock. Grey stars indicate significant change of a sample treated with FGFR inhibitor (BGJ398) compared to its respective control without inhibitor (as indicated by the grey lines). *P < 0.05; **P < 0.01; ***P < 0.001; ****P < 0.0001 (paired Student’s t-test).

### FGF2 regulates ECM production by mammary fibroblasts

Fibroblasts are main producers of ECM in most tissues. Therefore, we investigated the role of FGF signaling in mammary fibroblasts in ECM production. Because our previous tests showed that from the FGFs tested, FGF2 is the most potent regulator of multiple functions in mammary fibroblasts, we investigated only the effects of FGF2 on ECM production. To this end, fibroblasts were cultured in conditions that favor ECM production, at high confluence and with L-ascorbic acid supplemented in the medium. In these cultures, we analyzed expression of candidate genes involved in ECM production by qPCR, and the ECM produced by fibroblasts was investigated using chromogenic assays, immunofluorescence and scanning electron microscopy.

The qPCR analysis revealed that FGF2 decreased expression of collagen genes *Col1a1*, *Col1a2*, and *Col3a1*, upregulated *Col7a1*, but had no effect on *Col4a1* or *Col4a2* expression (Fig. 3D). Furthermore, FGF2 decreased expression of *Lox*, the enzyme involved in covalent cross-linking of collagen fibrils (Kagan and Li, 2003), and increased expression of *Mmp3* and *Timp1* (Fig. 3D) but no significant change in *Loxl3* (Fig. S3A) or *Mmp11* expression was detected (Fig. 3D). FGF2 also induced expression of fibronectin (*Fn1*), osteopontin (*Spp1*) and heparan sulfate proteoglycan (HSPG) syndecan 1 (*Sdc1*), but had no effect on expression of HSPG genes *Hspg2, Sdc2, Sdc3*, or *Sdc4* (Fig. 3D, Fig. S3A). Expression of periostin (*Postn*) and hyaluronan synthase genes *Has2* and *Has3* was downregulated by FGF2 (Fig. 3D). Furthermore, we also detected increased expression of transforming growth factor beta (*Tgfb*), a major regulator of ECM, and *Dusp6*, an FGFR-ERK signaling target gene (Fig. 3D). This pattern of gene expression, observed after 3 days of FGF2 treatment, was predominantly sustained also on day 7 of fibroblast ECM culture. However, *Postn* and *Sdc1* no longer showed significant change in gene expression, and *Acta2* and *Mmp11* showed significant downregulation (Fig. S4A).

Picrosirius red staining of fibroblast ECM cultures revealed presence of fibrillar collagen in the ECM (Fig. 4A). The amount of collagen in cultures treated with FGF2 was significantly lower than in cultures treated with serum or serum with FGFR inhibitor. When compared to cultures with no FGF, the FGF2-treated cultures appeared to contain similar amount of collagen (Fig. 4A, Fig. S5B). However, inspection of the ECM cultures revealed significantly lower density of cells in cultures with no FGF or with both FGF2 and BGJ398 than in cultures with FGF2 (Fig. S5A), which was consistent with the role of FGF2 in fibroblast proliferation and survival (Fig. 2A, Fig. S2A, B). When the amount of collagen detected in ECM cultures was normalized to the cell number, FGF2-treated cultures displayed significantly less collagen than cultures with no FGF or with FGF2 and BGJ398 (Fig. 4B), which was in concordance with decreased expression of fibrillar collagen genes in response to FGF2 detected by qPCR (Fig. 3D).

**Figure 4.**
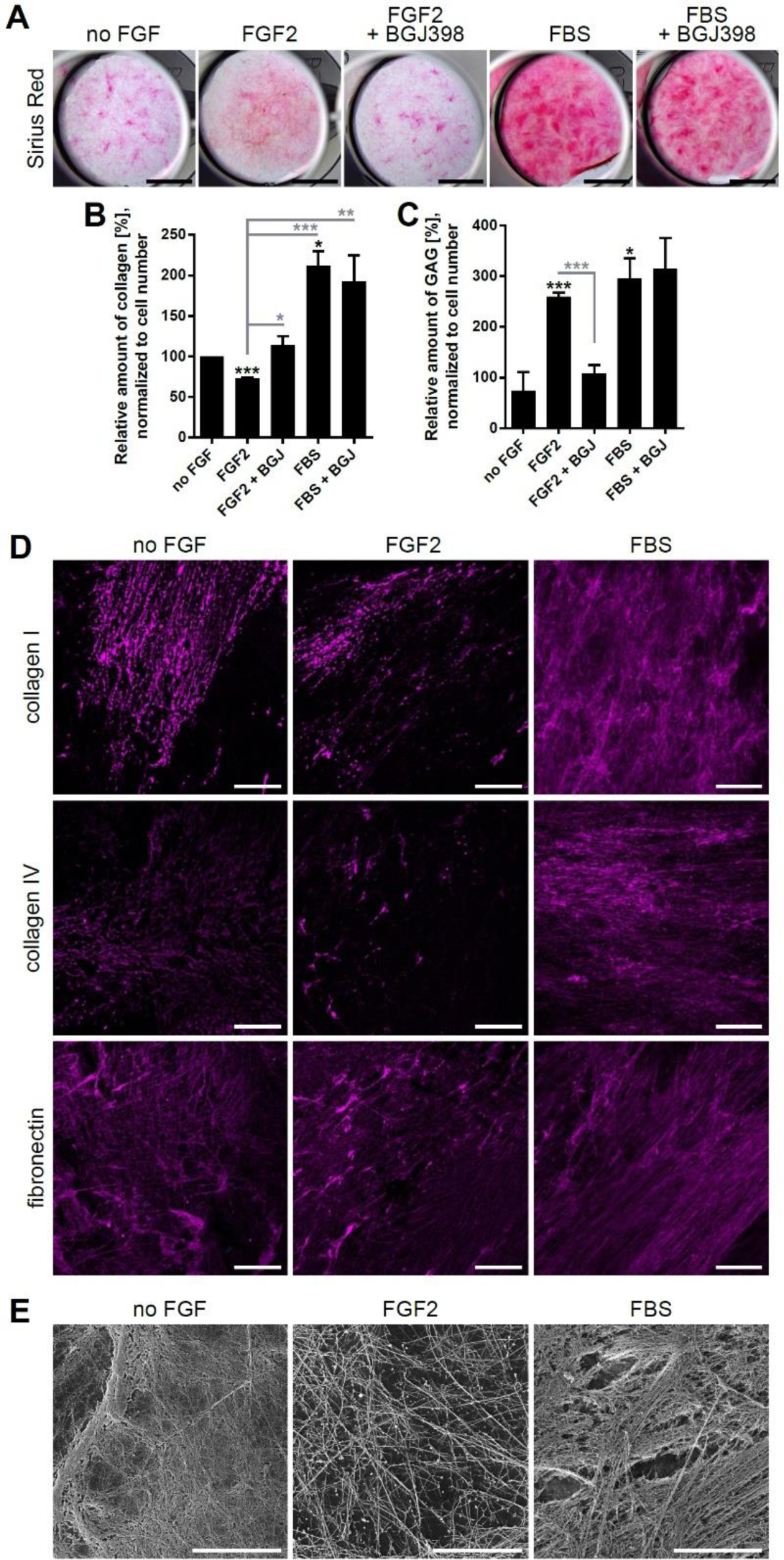
FGF2 regulates ECM production. (**A, B**) Detection of collagen production using Sirius Red. (**A**) Photographs of Sirius Red stained cell cultures. Scale bar, 5 mm. (**B**) Quantification of collagen amount. The graphs show the relative amount of collagen under different treatments in comparison to culture with no FGF, normalized to cell number, as mean + s.d.; n = 2-4. (**C**) Quantification of glycosaminoglycans (GAG). The graphs show the relative amount of GAG under different treatments in comparison to culture with no FGF, normalized to cell number, as mean + s.d.; n = 2-4. (**D**) Immunofluorescent staining of CDMs for collagen I, collagen IV and fibronectin. Scale bar, 20 μm. (**E**) Representative images from scanning electron microscopy of decellularized cell-derived matrices. Scale bar, 10 μm.

Analysis of glycosaminoglycans (GAGs) in the fibroblasts ECM cultures by Alcian Blue staining revealed significant upregulation of GAGs in FGF2-treated cultures in comparison to mock-or FGF2 and BGJ398-treated cultures (Fig. 4C, S5C). The amount of GAGs in serum-and serum with BGJ398-treated cultures was underestimated using this method. The bound dye inefficiently eluted from the complex ECM formed in serum-and serum with BGJ398-treated cultures.

Immunofluorescent (IF) staining for collagen I, collagen IV and fibronectin detected presence of these proteins in fibroblast-derived matrices and showed differences in protein amount and organization pattern (Fig. 4D). Overall, the matrices from serum-supplied cultures contained more collagen I, collagen IV and fibronectin and their staining patterns revealed more complex ECM meshwork than the matrices from serum-low cultures. The matrices from serum-low cultures showed low signal for collagen IV and the staining pattern was more focally organized in FGF2-treated matrices. Moreover, matrices from FGF2-treated cultures displayed higher signal for fibronectin than matrices from cultures with no FGF, and lower amount of collagen I than matrices from cultures with no FGF.

Scanning electron microscopy revealed structural differences in matrices produced by mammary fibroblasts. The most complex and highly organized matrix was produced by fibroblasts cultured with serum (Fig. 4E). Fibroblasts cultured in serum-low medium without FGF supplementation produced a less organized matrix of rather thin fibers and material of less distinctive structure. The matrix produced by fibroblasts treated with FGF2 contained thick fibers with small bulbous structures (Fig. 4E). This phenotype reflected the sum effect of FGF2-induced ECM gene expression changes and traction force-mediated remodeling.

### FGF2 signaling in fibroblasts enhances fibroblast-induced branching of mammary epithelium

Stromal fibroblasts are important regulators of mammary epithelial morphogenesis (Hammer et al., 2017; Koledova et al., 2016; Morsing et al., 2016; Peuhu et al., 2017). Therefore, we investigated the role of FGF2 signaling in fibroblasts in epithelial branching morphogenesis using 3D cultures of mammary organoids alone, fibroblasts alone, and co-cultures of organoids with fibroblasts. Using growth factor array, we detected low amount of FGF2 protein in organoid cultures, and increased amounts of FGF2 in fibroblast cultures and in organoid-fibroblast co-cultures (Fig. 5A, B). Another differentially abundant growth factor was amphiregulin (AREG), a mammary epithelium-derived protein (Ciarloni et al., 2007; Sternlicht et al., 2005). AREG was highly abundant in organoid cultures, barely detected in fibroblast cultures, and moderate amounts of AREG were found in organoid-fibroblast co-cultures (Fig. 5A, B). qPCR analysis of 3D (co-)cultures confirmed organoid-specific expression of *Areg* and fibroblast-specific expression of *Fgf2*, and revealed increased expression of *Fgf2* in co-cultures (Fig. 5C), suggesting organoid-induced *Fgf2* expression in fibroblasts. Moreover, we found that FGF2 treatment induced *Fgf2* expression in fibroblasts (Fig. S6A), suggesting an autocrine FGF2 signaling loop in fibroblasts.

**Figure 5.**
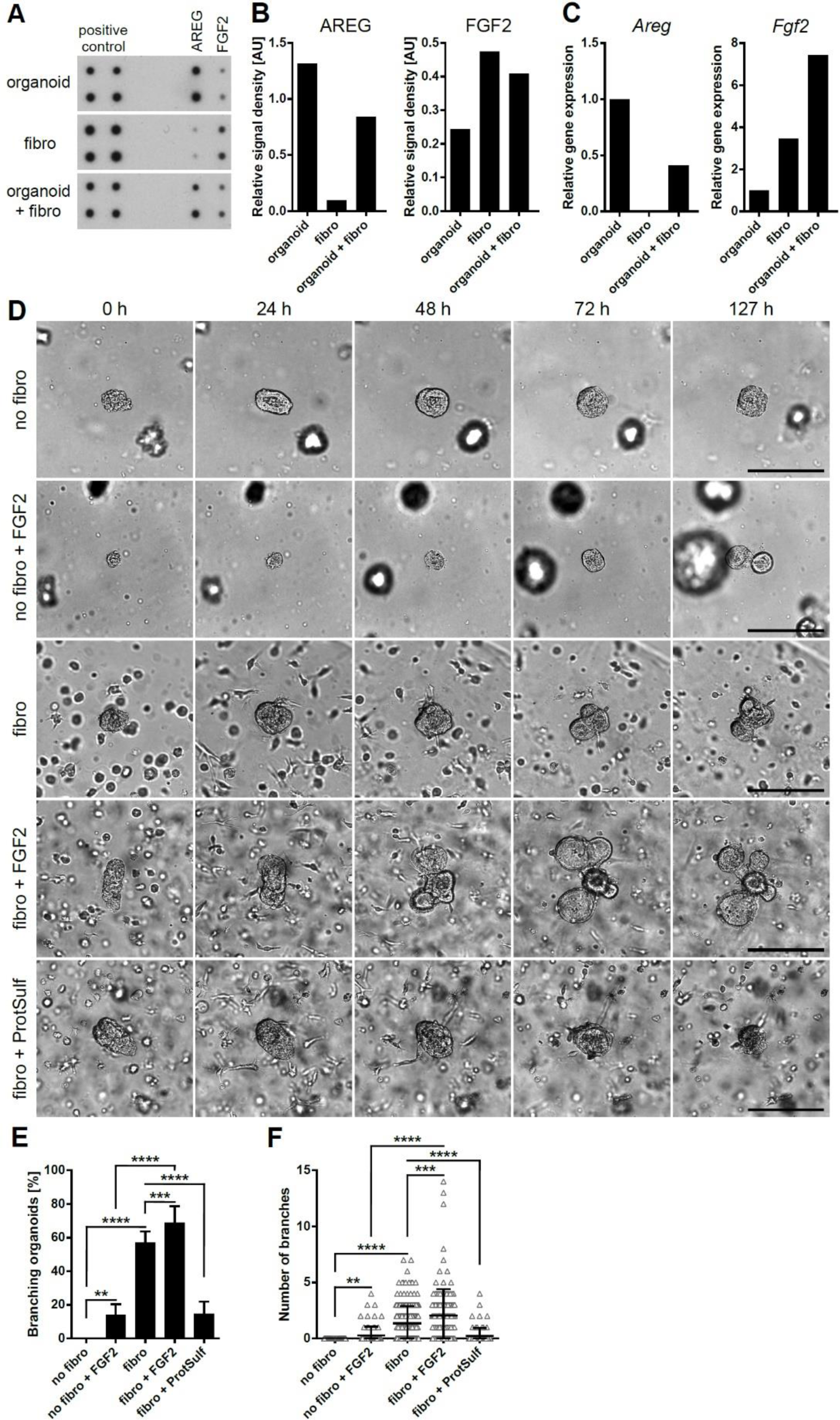
FGF2 signaling in mammary fibroblasts enhances fibroblast-induced branching of mammary epithelium. (**A, B**) Growth factor array analysis of 3D cultures of organoids, fibroblasts, and 3D co-cultures of organoids with fibroblasts. (**A**) Scans of growth factor arrays. (**B**) Quantification of signal for AREG and FGF2, normalized to positive control. (**C**) qPCR analysis of *Areg* and *Fgf2* expression in 3D (co-)cultures; n = 1. (**D-F**) Analysis of mammary epithelial organoid branching in response to FGF2 and/or fibroblasts. 3D cultures of organoids (no fibro) and co-cultures of organoids and fibroblasts (fibro) were treated with FGF2 or protamine sulfate (ProtSulf) as indicated and imaged for 5 days by time-lapse microscopy. **(D)** Representative snapshots from time-lapse microscopy. Scale bar, 100 μm. (**E, F**) Quantification of epithelial organoid branching. The plots show (**E**) percentage of branched organoids as mean + s.d. and (**F**) the number of branches per organoid (each triangle indicates one organoid); the lines show mean ± s.d., n = 4-10 (66 – 238 organoids). **P < 0.01; ***P < 0.001; ****P < 0.0001 (Student’s t-test).

Next, we analyzed the effect of fibroblasts and FGF2 on epithelial branching using time-lapse microscopy of 3D (co-)cultures. Mammary epithelial organoids did not branch in organoid only cultures in basal medium without FGF2. In co-cultures with fibroblasts, 57% of mammary organoids underwent branching (Fig. 5D-F). The frequency of fibroblast-induced branching was significantly increased (to 69%) by addition of 1 nM FGF2, while in organoid only cultures, 1 nM FGF2 induced branching only at low frequency (14%) (Fig. 5D-F). Similarly, the highest number of branches per organoid (Fig. 5F) and the highest number of branches per branched organoid (Fig. S6B) were formed in co-cultures with fibroblasts and 1 nM FGF2. Furthermore, in co-cultures with fibroblasts and 1 nM FGF2, the organoid branches were the most prominently developed. Importantly, inhibition of paracrine and autocrine FGF signaling by protamine sulfate, an inhibitor of FGF co-receptor heparan sulfate (Wolzt et al., 1995), abrogated fibroblast-induced branching (Fig. 5D-F, Fig. S6B). Taken together, our results provide evidence of a role for FGF signaling in fibroblast-induced branching morphogenesis.

## Discussion

Fibroblasts are important regulators of mammary epithelial morphogenesis (Hammer et al., 2017; Koledova et al., 2016; Kuperwasser et al., 2004; Morsing et al., 2016; Peuhu et al., 2017; Zhang et al., 2002), but the regulation of fibroblast function is incompletely understood. In this study, we identified functional components of FGF signaling in mammary fibroblasts and their roles in regulation of multiple biological functions, including fibroblast proliferation, migration, ECM production and remodeling, and interactions with mammary epithelium during mammary epithelial branching morphogenesis.

We found expression of FGFR1 and FGFR2 in the mammary fibroblasts, both in their IIIc variants. This is consistent with the general observation of the distinct spatial expression patterns of the FGFR splice variants that the IIIc variants are preferentially found in the mesenchyme, while the IIIb variants are more common in epithelia (Kettunen et al., 1998; Orr-Urtreger et al., 1993; Rice et al., 2003). Moreover, in agreement with the FGF ligand specificity of the IIIc variants of FGFR1 and FGFR2 (Ornitz and Itoh, 2015), we detected functional response (such as proliferation and 2D migration) of mammary fibroblasts to FGF2 and FGF9, but no functional response was elicited by FGF7 or FGF10, which activate specifically FGFR IIIb variants. The lack of functional outcome in response to FGF7 and FGF10 corresponded to only weak and transient activation of ERK1/2, while FGF2 and FGF9 elicited strong and sustained ERK1/2 activation. These observations are in agreement with previous study that ERK1/2 signaling dynamics is the key determinant of cellular response to FGF signaling (Zhu et al., 2010). Moreover, differences in AKT signaling engagement in response to FGF2 and FGF9 that we observed, and/or distinct FGFR signaling kinetics (Francavilla et al., 2013) could underlie the distinct capability of FGF2 to promote fibroblast migration in 3D ECM and force-mediated collagen remodeling.

Our investigation of the mechanism of FGF2-induced fibroblast migration in 3D ECM by inhibition of candidate molecules revealed requirement for Rho GTPases (Rac1, Cdc42), which regulate actin dynamics, and MLCK, which phosphorylates myosin regulatory light chain 2 (MLC2) to regulate activity and assembly of myosin II filaments (Sit and Manser, 2011). Intriguingly, inhibition of ROCK, which also phosphorylates MLC2, enhanced fibroblast migration. This could be, at least in part, explained by MLCK and ROCK acting on distinct myosin II pools (Totsukawa et al., 2004). Inhibition of myosin II using blebbistatin induced massive amoeboid-like migration of fibroblasts, which is consistent with a role for low contractility in this switch in migration mode (Liu et al., 2015).

ECM signals, including ECM composition, stiffness and topology, regulate mammary epithelial cell adhesion, migration, proliferation, apoptosis, survival and differentiation, and thereby determine mammary epithelial morphogenesis (Bonnans et al., 2014; Schedin and Keely, 2011). For example, patterning of collagen fibers in the mammary gland stroma regulates the orientation of TEBs during branching morphogenesis (Brownfield et al., 2013). Recent publications reported important role of fibroblasts in regulation of ECM composition, abundance and organization, and thereby in regulation of mammary epithelial branching morphogenesis (Feinberg et al., 2018; Hammer et al., 2017; Koledova et al., 2016; Morsing et al., 2016; Peuhu et al., 2017). We found that FGF signaling in fibroblasts regulates expression of several ECM genes, including collagens. FGF2 downregulated expression of genes for fibrillar collagens I and III, the major structural proteins of connective tissues, including mammary gland ECM (Goddard et al., 2016). An opposite trend showed FGF2 in regulation of collagen VII, an anchoring fibril collagen that binds type I and type III collagen and is crucial for the function and stability of ECM (Nyström et al., 2013). FGF2 was also found to regulate expression of several genes involved in regulation of collagen abundance and organization, including *Lox*, *Mmp3*, and *Timp1*, a regulator of MMPs. Moreover, FGF2 induced expression of *Tgfb*, a growth factor involved in regulation of ECM production and remodeling during development as well as cancer progression (Moses and Barcellos-Hoff, 2011). Furthermore, Alcian blue staining revealed upregulation of GAGs in the fibroblasts-derived ECM by FGF2, what corresponded to upregulation of heparan sulfate proteoglycan (HSPG) *Sdc1*. Because HSPGs bind FGFs and act as FGF signaling co-receptors (Bernfield et al., 1999; Mundhenke et al., 2002), such regulation could form a positive FGF signaling loop in mammary fibroblasts.

Mammary fibroblasts reorganize collagen fibers by exerting traction forces during cell adhesion and migration in 3D ECM (Peuhu et al., 2017). We found that FGF signaling promotes traction force-mediated collagen remodeling. Activity of MMPs or LOX, the proteins involved in collagen remodeling by proteolytic cleavage or crosslinking, respectively (Bonnans et al., 2014), did not contribute to FGF2-induced collagen remodeling in collagen contraction assay. Moreover, although FGF2 regulates fibroblast proliferation, too, differences in cell proliferation in response to FGF ligands were unlikely to account for the distinctive FGF2-induced collagen gel contraction because inhibition of cell proliferation did not abrogate it. This is consistent with the reports that fibroblasts do not proliferate in floating collagen gels (Grinnell, 2000).

Furthermore, fibroblasts orchestrate mammary epithelial morphogenesis using paracrine signaling (Koledova et al., 2016; Zhang et al., 2002). In this study we showed that fibroblasts produce FGF2 and that co-culture of fibroblast with mammary epithelial organoids induced organoid branching. Moreover, we showed that *Fgf2* expression is positively regulated by FGF signaling in fibroblasts and that stimulation of FGF2 production by fibroblasts by low dose FGF2 increased epithelial branching frequency. FGF2 is a potent inducer of mammary epithelial branching (Ewald et al., 2008) in a dose-dependent manner (Zhang et al., 2014a). In the co-cultures treated with FGF2, the epithelium was unavoidably exposed to FGF2, too. But the dose that we used to stimulate fibroblasts was by itself inefficient in induction of pronounced epithelial branching. Therefore, we conclude that the epithelial branching in FGF2-treated co-cultures was predominantly caused by fibroblast-mediated signaling. Moreover, inhibition of FGF-FGFR signaling complex assembly by inhibitor of heparan sulfate abrogated fibroblast-induced branching, further supporting the role of fibroblast-produced FGF2 in epithelial branching.

In the mammary gland, and analogously in the 3D co-culture, the communication between stromal and epithelial cells is bidirectional, i.e. epithelial cells signal to stromal cells, too. AREG is one of the known epithelium-derived signal for stromal cells required for epithelial branching (Ciarloni et al., 2007; Sternlicht et al., 2005). In our experiments, we detected AREG expression by organoids and moderate levels of AREG were present in the co-cultures. However, it remains to be determined how FGF2 expression in fibroblasts is regulated and whether AREG plays role in it.

Together, we presented evidence for a role of FGF signaling in mammary fibroblasts in mammary epithelial branching morphogenesis by regulation of ECM remodeling and paracrine signaling. Our findings from 2D and 3D cultures should be further tested in the context of the whole mammary gland using mouse models, which would provide physiologic, more complex microenvironment and allow for analysis of long-term effects of FGF signaling modulation in fibroblasts. Moreover, fibroblasts are a key component of breast cancer stroma and a determinant of breast cancer progression (Ahn et al., 2012; Cid et al., 2018; Kalluri and Zeisberg, 2006). Their protumorigenic effects include production of paracrine proliferative and proinvasive signals to cancer cells (Bernard et al., 2018; Orimo et al., 2005), ECM remodeling to enable invasion of cancer cells (Gaggioli et al., 2007), and enabling immune evasion (Chakravarthy et al., 2018). Because increased expression of FGF ligands, including FGF1, FGF2 and FGF7, was found in breast cancer stroma in comparison to normal breast stroma (Finak et al., 2008; Relf et al., 1997), it is relevant to investigate the role of FGF signaling in breast cancer fibroblasts.

## Materials and Methods

### Primary mammary fibroblast isolation and culture

Primary mammary fibroblasts were isolated from 6-8 weeks old female ICR mice as previously described (Koledova, 2017). Briefly, the mice were euthanized by cervical dislocation according to protocol approved by veterinary committee and in compliance with the Czech and European laws for the use of animals in research. The mammary glands were removed, mechanically disintegrated and partially digested in a solution of collagenase and trypsin [2 mg/ml collagenase A, 2 mg/ml trypsin, 5 μg/ml insulin, 50 μg/ml gentamicin (all Sigma/Merck), 5% fetal bovine serum (FBS; Hyclone/GE Healthcare) in DMEM/F12 (Thermo Fisher Scientific)] for 30 min at 37°C. Resulting tissue suspension was treated with DNase (20 U/ml; Sigma/Merck) and exposed to five rounds of differential centrifugation, which resulted in separation of epithelial and stromal fractions. The cells from stromal fraction were pelleted by centrifugation, suspended in fibroblast cultivation medium (10% FBS, 1× ITS, 100 U/ml of penicillin, and 100 μg/ml of streptomycin in DMEM) and incubated on cell culture dishes at 37°C, 5% CO_2_ for 30 min. Next, selection by differential attachment was performed to remove non-fibroblast cells: After the 30 min incubation, unattached cells were washed away, the cell culture dishes were washed with PBS and fresh fibroblast medium was provided for the cells. The cells were cultured until about 80% confluence. During the first cell subculture by trypsinization, a second round of selection by differential attachment was performed, when the cells were allowed to attach only for 15 min at 37°C and 5% CO_2_. The cells were expanded and used for the experiments until passage 5.

### Immunohistochemistry of mammary gland

Mammary glands from 6-week-old female ICR mice were fixed with neutral buffered formalin overnight at room temperature (RT). Next day the tissue was process via standard procedure for paraffin embedding. Paraffin sections were cut (5 μm thickness), deparaffinized using xylene and rehydrated. Antigens were retrieved using Tris-EDTA buffer, pH 9 (Dako) and endogenous peroxidase activity was blocked using 3% hydrogen peroxide. The sections were blocked in PBS with 10% FBS and incubated with primary antibody (Table S1) for 1 h at RT. After washing, sections were incubated with secondary antibody (anti-mouse, EnVision+ Dual Link System-HRP; Dako) for 30 min at RT. After washing, bound secondary antibody was detected using Liquid DAB+ Substrate Chromogen System (Dako). The nuclei were stained with Mayer’s hematoxylin, dehydrated and mounted in Pertex (Histolab Products). The samples were photographed using Leica DM5000 equipped with Leica DFC480 camera.

### Immunofluorescence staining of fibroblasts

Fibroblasts were cultured on coverslips, fixed with neutral buffered formalin, permeabilized with 0.05% Triton X-100 in PBS and blocked with PBS with 10% FBS. Then the cells were incubated with primary antibodies (Table S1) for 2 h at RT. After washing, the cells were incubated with secondary antibodies (Table S1) and Alexa Fluor 488 Phalloidin (Thermo Fisher Scientific). Then the cells were washed, stained with DAPI (1 μg/ml; Merck) for 10 min and mounted in Mowiol (Merck). The cells were photographed using Eclipse Ti microscope (Nikon).

### MTT assay

For FGF proliferation assay, 5000 or 7000 cells per well were seeded in 96-well plates in fibroblast cultivation medium. The next day the plates were carefully washed three times with PBS and supplied with serum-low medium (0.2% FBS, 0.1× ITS, 100 U/ml of penicillin, 100 μg/ml of streptomycin in DMEM) and incubated for 24 h at 37°C, 5% CO_2_. The next day FGF ligands (to 5 nM) and heparin (to 2 μg/ml; Merck) were added. All treatments were done in intraexperimental triplicates. The cells were incubated with FGF ligands for 24 h. Then MTT (Merck) was added to the plate to the final concentration 0.45 mg/ml and the plates were incubated for 5 h at 37°C, 5% CO_2_. Then medium was aspirated and formazan crystals were dissolved in 10% SDS, 0.01 M HCl. MTT absorbance (at 570 nm) was measured using Synergy HTX microplate reader.

### Scratch assay

Fibroblasts were seeded in 12 well plate and grown to full confluence in fibroblast cultivation medium. Then the scratch was induced using a 200 μl pipette tip, the cells were washed three times with PBS and treated with 5 nM FGF2, FGF7, FGF9, or FGF10 in serum-low medium (0.2% heat-inactivated FBS [FBS-HI], 0.1× ITS, 100 U/ml of penicillin, and 100 μg/ml of streptomycin in DMEM) with 2 μg/ml heparin. The plate was incubated in humidified atmosphere of 5% CO_2_ at 37°C on Olympus IX81 microscope equipped with Hamamatsu camera and CellR system for time-lapse imaging. The scratch areas (6 non-overlapping areas per well) were photographed every hour for 24 h or until they were closed by migrating cells (when cells from opposite sides of scratch meet) in at least one of imaged positions. The first occasion of scratch closure was set as experimental endpoint. The images were exported and analyzed using Image J (NIH). The first and endpoint image for each position were overlaid, cells that migrated were counted and their migration distance was measured (perpendicular to scratch median line). Average total migration distance per condition was calculated and compared to average total migration distance of cells with no FGF treatment.

### Transwell migration assay

1×10^5^ fibroblasts were seeded in the upper chamber of transwell (polycarbonate membrane, 8 μm pore size; Corning) in transwell assay medium (0.1% BSA, 2 μg/ml heparin in DMEM). In the lower chamber, transwell assay medium with 5 nM FGF2, FGF7, FGF9, or FGF10 was added. The cells were incubated for 10-11 h at 37°C, 5% CO_2_. Then the filter side of the upper chamber was wiped with a cotton swab to remove non-migratory cells and the remaining cells on the opposite side of the membrane were fixed with 4% paraformaldehyde for 15 min, washed with distilled water and stained with 0.1% crystal violet (Merck) for 15 min. After washing with distilled water, the membranes were cut from the inserts and the number of migrated cells was counted using Olympus IX81.

### Fibrosphere 3D migration assay

The fibroblasts were aggregated into fibrospheres by overnight culture at high density in fibroblast medium in bacterial dishes as previously described (Koledova, 2017). The fibrospheres were collected, washed three times with PBS, mixed with Matrigel (Corning) and plated in 3D domes. The cultures were incubated for 45 min at 37°C, 5% CO_2_, then basal fibrosphere medium (1× ITS, 100 U/ml of penicillin, and 100 μg/ml of streptomycin in DMEM) was added, supplied with 5 nM FGF ligands and/or inhibitors (Table S2). The cultures were incubated in humidified atmosphere of 5% CO_2_ at 37°C on Olympus IX81 microscope equipped with Hamamatsu camera and CellR system for time-lapse imaging. The fibrospheres were photographed every 60 min for 48 h. The images were exported and analyzed using Image J. For each fibrosphere, length of all protrusions was measured radially from the edge of the fibrosphere body to the end of the protrusion.

### Analysis of FGF signaling dynamics by Western blotting

Fibroblasts were grown to desired confluence, washed three times with PBS and serum-starved in DMEM with 0.05× ITS, 100 U/ml of penicillin, and 100 μg/ml of streptomycin. Next day the cells were treated with 5 nM FGF ligands in DMEM with 4 μg/ml heparin for 5 to 60 min. After treatment, cells were immediately washed twice with ice-cold PBS and lysed in the RIPA buffer [150 mM NaCl, 1.0% NP-40, 0.5% sodium deoxycholate, 0.1% SDS, 50 mM Tris, pH 8.0] supplied with proteinase and phosphatase inhibitors (10 mM β-glycerophosphate, 5 mM NaF, 1 mM Na_3_VO_4_, 1 mM dithiotreitol, 0.5 mM phenylmethanesulphonylfluoride, 2 µg/ml aprotinin, 10 µg/ml leupeptin; all Merck). Protein lysates were homogenized by vortexing, cleared by centrifugation and protein concentration was measured using the Bradford reagent. Denatured, reduced samples were resolved on 10% SDS-PAGE gels and blotted onto PVDF membranes (Merck). Membranes were blocked with 5% non-fat milk in PBS with 0.05% Tween-20 (Merck; blocking buffer) and incubated with primary antibodies (Table S1) diluted in blocking buffer overnight at 4°C. After washing in PBS with 0.05% Tween-20, membranes were incubated with horseradish peroxidase-conjugated secondary antibodies (Table S1) for 1 h at RT. Signal was developed using an ECL substrate (100 mM Tris-HCl, pH 8.5, 0.2 mM coumaric acid, 1.25 mM luminol, 0.01% H_2_O_2_) and exposed on X-ray films (Agfa), which were then scanned and band density was analyzed using ImageJ. Phosphorylated and total proteins and actin were analyzed on a single blot.

### Analysis of ECM production

Fibroblasts were cultured to full confluence, then washed three times with PBS and treated with ECM starvation medium (1% FBS-HI, 1× ITS, 100 U/ml of penicillin, and 100 μg/ml of streptomycin in DMEM) supplemented with 50 μg/ml L-ascorbic acid and 1 nM FGF2, 1 nM FGF2 and 0.1 μM BGJ398, 0.1 μM BGJ398, 10% FBS, or 10% FBS and 0.5 μM BGJ398. All treatments were done in intraexperimental duplicates or triplicates. The cells were cultured for 3 and 7 days. Fresh L-ascorbic acid was added to the cultures every 2 days. Fresh medium was added to the cultures after 3 days.

For detection of fibrous collagen, the fibroblast cultures were washed with PBS, fixed with Bouin’s solution (75 ml of saturated picric acid solution, 25 ml of 40% formaldehyde, 5 ml of glacial acetic acid) for 1 h, washed with tap water and incubated for 1.5 h in 0.1% (w/v) Sirius red (Merck) in saturated picric acid solution with mild shaking. After discarding the stain, the cell cultures were washed with 0.01 M HCl to remove unbound dye, dried and photographed using stereo microscope Leica M165 FC equipped with camera Leica DFC450C. Subsequently the bound dye was eluted using 0.1 M NaOH, extracts were collected, clarified by centrifugation and their absorbance at 550 nm was measured using Synergy HTX reader (BioTek).

For detection of glycosaminoglycans, the fibroblast cultures were washed with PBS and fixed with 4% formaldehyde (Bio-Optica) for 1 h at 4°C. After washing with 0.1 M HCl, the cultures were incubated with 0.5% Alcian blue 8GX (Serva) in 0.1 M HCl. Afterwards the staining solution was discarded and the cultures were extensively washed with deionized water to remove any unbound stain. The cultures were scraped into 1% SDS to extract bound dye, centrifuged and the absorbance of the supernatant at 350 nm was measured using Synergy HTX reader.

For quantification of cell number, the medium was removed and fibroblast cultures were supplied with 10 µg/ml resazurin (Merck) in DMEM and the plates were incubated for 3 h at 37°C, 5% CO_2_. Resorufin fluorescence (excitation at 560 nm, emission at 590 nm) was measured using Synergy H4 Hybrid multi-mode microplate reader (BioTek).

### Production of cell-derived matrices

Fibroblasts were cultured on gelatin-coated coverslips to full confluence, then washed three times with PBS and treated with ECM starvation medium (1% FBS-HI, 1× ITS, 100 U/ml of penicillin, and 100 μg/ml of streptomycin in DMEM) with no FGF, 1 nM FGF2, or 10% FBS, and supplemented with 50 μg/ml L-ascorbic acid (Merck). The cells were cultured for 11 days and the medium was changed every day. After 11 days of cultivation, the cultures were decellularized according to published protocol (Kaukonen et al., 2017) and processed for scanning electron microscopy.

### Scanning electron microscopy

The samples were fixed with 3% glutaraldehyde in 100 mM sodium cacodylate buffer, pH 7.4 for 45 min, washed with cacodylate buffer and dehydrated in ethanol series. Samples were dried in Critical Point Dryer CPD 030 (Bal-Tek) and then sputter-coated with gold in SCD 040 (Balzers Union) at 30 mA for 3 min. Gilded specimens were analyzed with scanning electron microscope VEGA TS 5136 XM (TESCAN).

### Real-time quantitative PCR (qPCR)

RNA from fibroblasts was isolated using RNeasy Mini Kit (Qiagen) according to the manufacturer’s instruction. RNA concentration was measured using NanoDrop 2000 (Thermo Fisher Scientific). RNA was transcribed into cDNA by using Transcriptor First Strand cDNA Synthesis Kit (Roche) or TaqMan Reverse Transcription kit (Life Technologies). Real-time qPCR was performed using 5 ng cDNA, 5 pmol of the forward and reverse gene-specific primers each (primer sequences are shown in Table S3) in Light Cycler SYBR Green I Master mix (Roche) on LightCycler 480 II (Roche). Relative gene expression was calculated using the ΔΔCt method and normalization to two housekeeping genes, β-actin (*Actb*) and Eukaryotic elongation factor 1 γ (*Eef1g*).

### Collagen contraction assay

Fibroblasts were collected from cell culture dishes by trypsinization, washed 3 times with PBS to remove any traces of serum and suspended in DMEM. Neutralized collagen was prepared by combing 12.5 volumes of collagen type I (Corning) with 1 volume of 0.22 M NaOH, 5× collagen reconstitution buffer (5× MEM, 20 μg/ml NaHCO_3_, 0.1 M Hepes), DMEM and fibroblast suspension to the final concentration 2.58 mg/ml collagen, 1× MEM, and final cell density 2.8×10^5^ cell/ml. Equal volumes of the collagen-fibroblast mixture were plated in 24-well BSA-coated plate (712 μl/well) and incubated at 37°C, 5% CO_2_ for 1 h before medium (1× ITS, 100 U/ml of penicillin, and 100 μg/ml of streptomycin in DMEM) with 5 nM FGF ligands and/or inhibitors, as required for the experiment, was added on the top. The gels were cultured 2 days and fixed using neutral buffered formalin. Fixed gels were photographed using Leica M165 FC equipped with camera Leica DFC450C. The images were merged using Adobe Photoshop and the gel area was measured in Image J. The extent of gel contraction was calculated as the percentage of the area of a non-contracted gel with no cells.

### 3D culture of mammary organoids and fibroblasts

3D culture of mammary organoids and fibroblasts was performed as previously described (Koledova and Lu, 2017). Briefly, the freshly isolated mammary organoids were embedded in Matrigel either alone or with mammary fibroblasts and plated in domes. After setting the gel for 45-60 min at 37°C, the cultures were overlaid with basal organoid medium (1× ITS, 100 U/ml of penicillin, and 100 μg/ml of streptomycin in in DMEM/F12), supplied with 1 nM FGF2 (Peprotech) or 50 μg/ml protamine sulfate (Merck) according to the experiment. The cultures were incubated in humidified atmosphere of 5% CO_2_ at 37°C on Olympus IX81 microscope equipped with Hamamatsu camera and CellR system for time-lapse imaging. The organoids were photographed every 60 min for 5 days with manual refocusing every day. The images were exported and analyzed using Image J. Organoid branching was evaluated from videos and it was defined as formation of a new bud/branch from the organoid. Organoids that fused with another organoid or collapsed after attachment to the bottom of the dish were excluded from the quantification.

Growth factor production in the 3D (co-)cultures was analyzed on day 3 from the media using Mouse growth factor array (Ray Biotech) and from cell lysates using qPCR. For qPCR analysis of *Fgf2* expression in response to FGF2 in fibroblasts, the fibroblasts were cultured in 3D for 24 h in serum-starvation medium (1× ITS, 0.2% FBS, 100 U/ml of penicillin, and 100 μg/ml of streptomycin in DMEM) and then treated with 5 nM FGF2 and 4 μg/ml heparin in serum-starvation medium for 24 h.

### Statistical analysis

Statistical analysis was performed using GraphPad Prism software using Student’s t-test. *P < 0.05, **P < 0.01, ***P < 0.001, ****P < 0.0001. Line plots and bar graphs were generated by GraphPad Prism and show mean ± standard deviation (s.d.). The number of independent biological replicates is indicated as n.

## Acknowledgements

We thank Julia Smelkova, Zuzana Garlikova, Michaela Klouckova, Katarina Mareckova and Anas Rabata for technical assistance and Jakub Cernek for help with image analysis.

## Funding

This study was supported by Czech Science Foundation [GJ16-20031Y to Z.K.]; and by funds from the Faculty of Medicine MU to junior researcher Z.K.

## Declaration of Interests

The authors declare that they have no competing interests.

